## Supplemental Figures 1-13 for "Self-organization of modular activity in immature cortical networks"

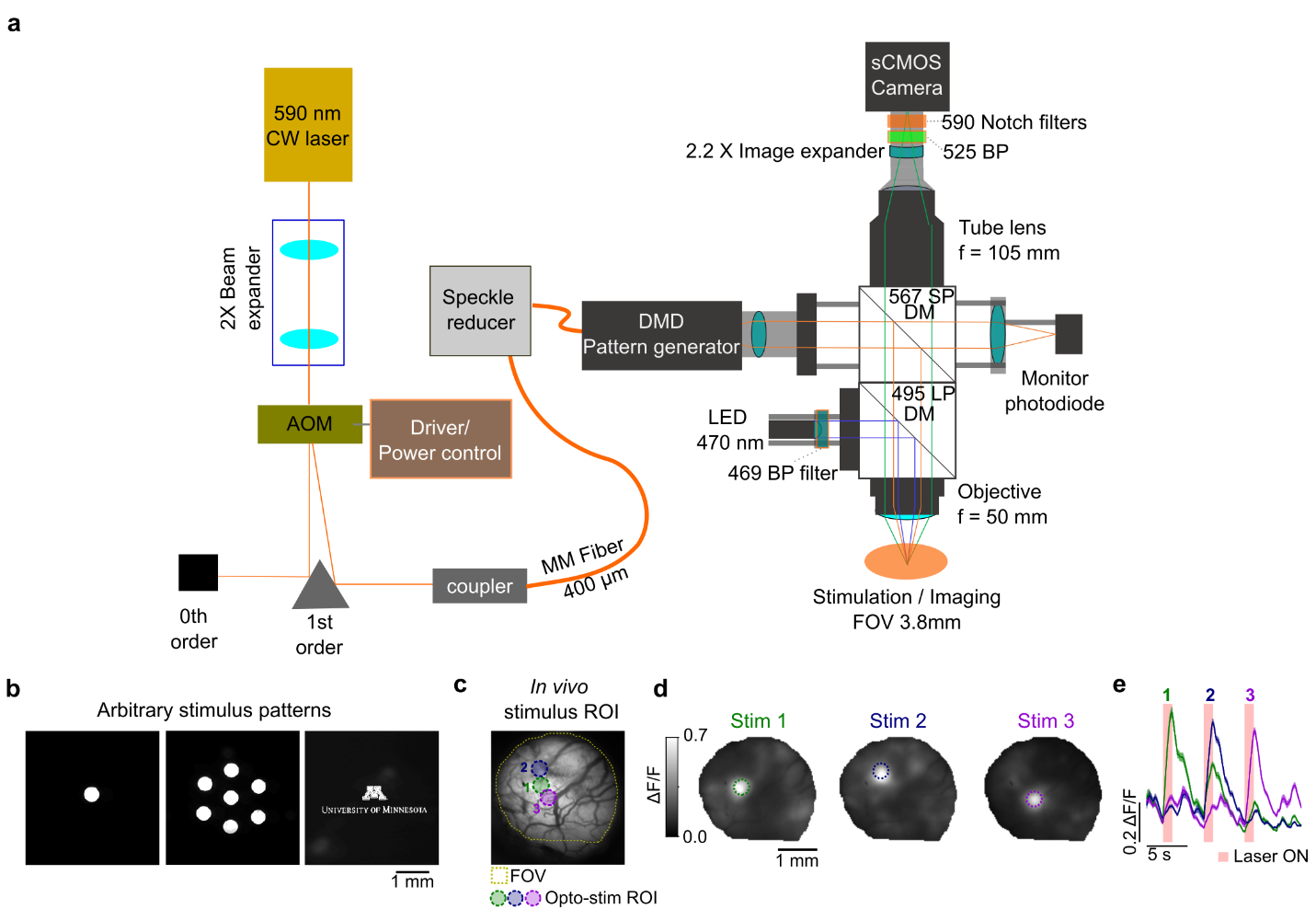

Fig. S1. All-optical stimulation and imaging of columnar cortical activity. (a) Optomechanical light path for mesoscopic simultaneous calcium imaging and optogenetic stimulation. For optogenetic stimulation, a 590 nm continuous wave (CW) laser controlled by an acousto-optic modulator (AOM) is coupled into a digital micromirror device (DMD) to project spatially defined patterns into the surface of the imaging plane. Stimulation artifacts are blocked from reaching the imaging camera with a 590 nm notch filter. (DM=dichroic mirror, SP, LP, BP=short-, long-, and band-pass filter, respectively, f=focal length) (b) Projection of stimulus patterns onto imaging plane, imaged by camera (with 590 nm notch filter removed to allow visualization of stimulus pattern). (c) Three example stimulus ROIs. (d-e) Optogenetic stimulation drives spatially specific responses. (d) Response at stimulus offset, mean across 20 trials. (e) Stimulus triggered average response for pixels within each stimulus ROI shown in (d) (mean, +/- SEM).

**a**

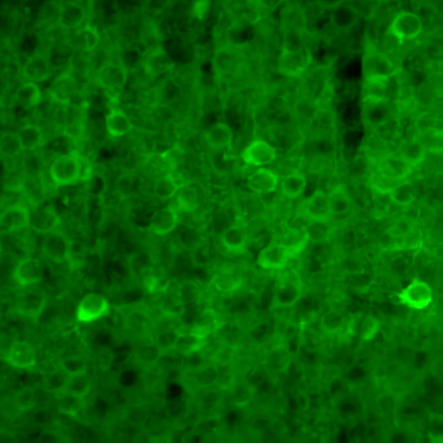

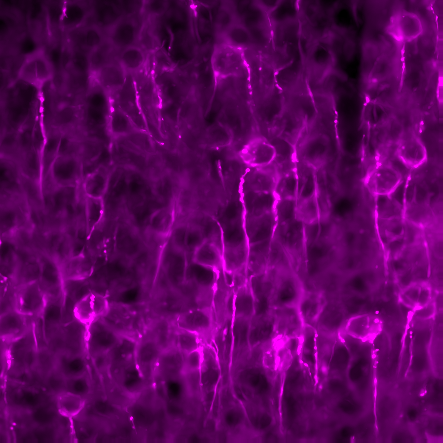

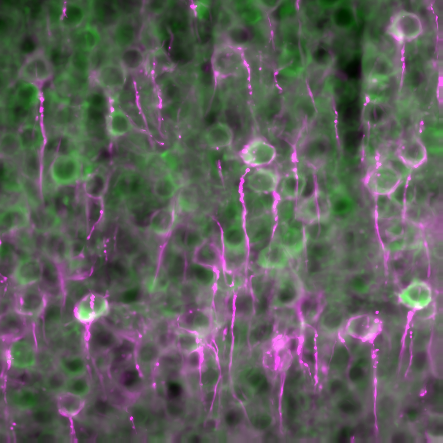

Fig. S2. Co-expression of GCaMP6s and Chrimson-ST in developing ferret visual cortex. (a) Confocal images of GCaMP6s (left, green), somatically tagged Chrimson (middle, magenta) in layer 2/3 of developing ferret visual cortex (P27). (Right) Merged GCaMP6s and Chrimson images.

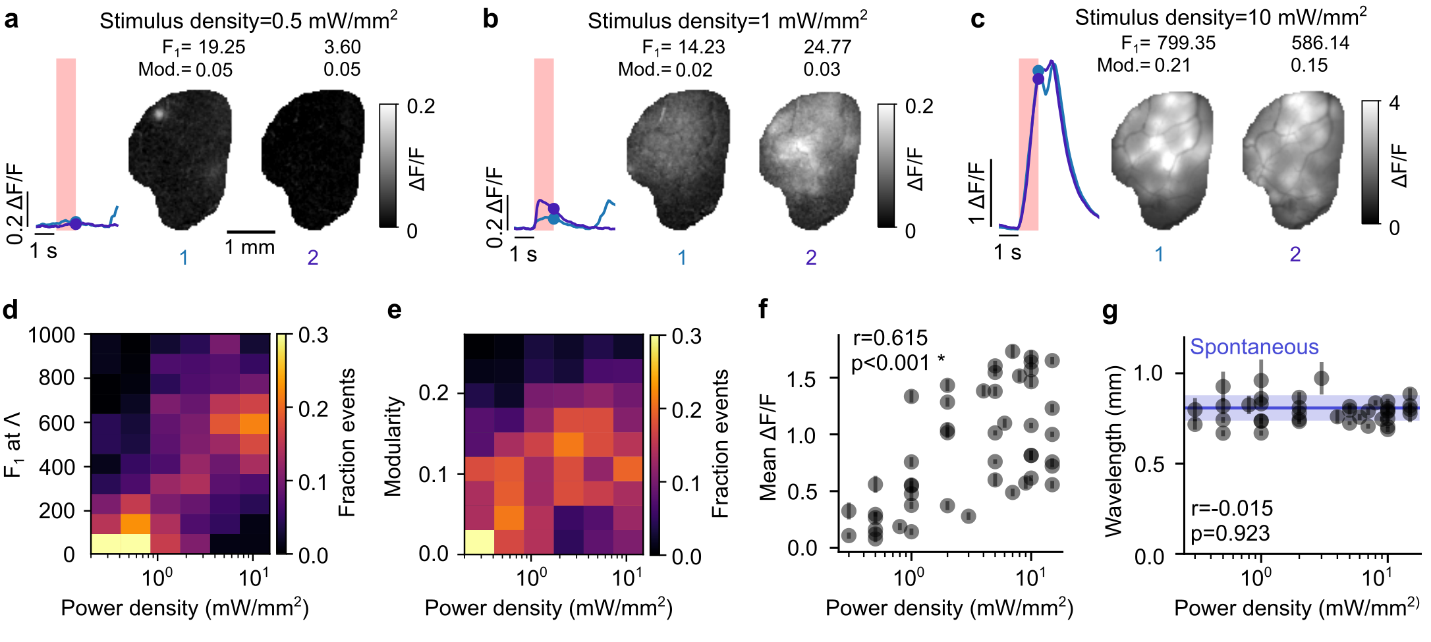

Fig. S3. Opto-evoked modularity increases nonlinearly with stimulus power, while the wavelength remains constant. (a-c) Uniform opto-evoked responses for low (0.5 mW/mm^2^, a), intermediate (1 mW/mm^2^, b) and high (10 mW/mm^2^, c) stimulus power. (*Left)* example traces of mean ∆F/F over ROI for 2 trials shown to right. (*Right*) Example opto-evoked activity at stimulus offset. Quantification of modular amplitude at the characteristic wavelength (F_1_), see Methods) and modularity (Mod.) are given for each event. (d) Modular amplitude of activity switches from a distribution peaked at low amplitude responses for low power densities to a distribution peaked at high amplitude responses at high power densities, consistent with the presence of a dynamic instability for sufficiently strong input drive. Density plot across animals, normalized within column. (e) Modularity shows non-linear increase with stimulus power. (f) Amplitude of overall activity (mean ΔF/F across field of view) increases with increasing power. (Mean (+/- SEM) of n=40 trials for each power density.) (g) The wavelength of opto-evoked events is invariant to power density (Pearson’s r=-0.015, p=0.923) and resides in a narrow band consistent with the wavelength of spontaneous activity (blue, mean +/- std. dev. across animals). Average wavelength (+/- SEM) across modular trials (trials with modularity >2 standard deviations over mean at baseline in 1 second window before stimulus onset) for each power density tested. For (D-G) data is pooled across 7 animals.

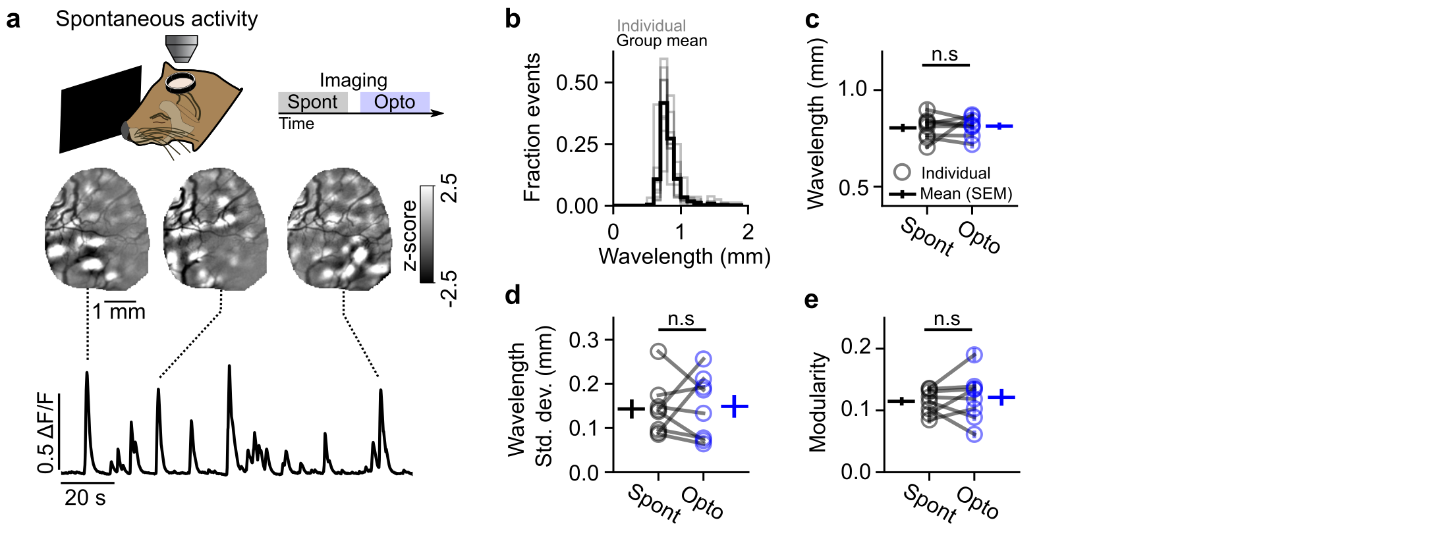

Fig. S4. Uniform opto-evoked activity has similar wavelength and modularity to spontaneous activity. (a) Experimental schematic. Spontaneous activity is highly modular, consistent with prior work. (b) Distribution of wavelength across spontaneous events is narrow, consistent with a characteristic wavelength (transparent lines=distribution in individual animals, opaque line=group mean). (c-e) Opto-evoked activity patterns and spontaneous activity share similar spatial wavelength (c), narrowness of wavelength distribution across patterns (d), and degree of modularity (e). (Open circles=average across individual animals, horizontal line=group mean +/- SEM.)

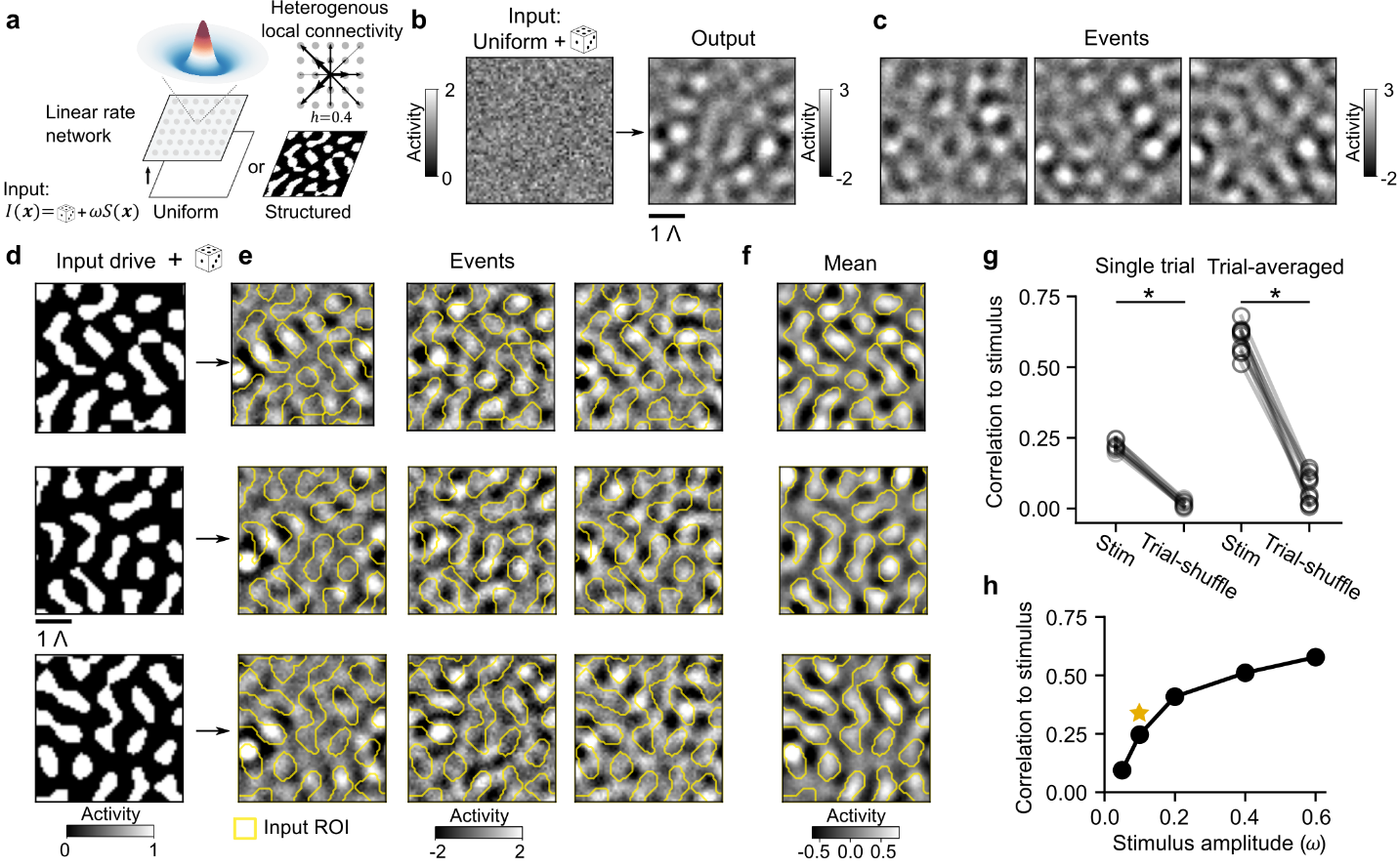

Fig. S5. Computational model of cortical network to study the predictions in Fig. 1c and d and the effect of noise. (a) Schematic of a linear rate model network with short-range interactions of local excitation and lateral inhibition (see Methods). Network connections are slightly heterogeneous, in line with models of spontaneous cortical activity (Smith et al. 2018) (See Fig. S10 for the homogenous case). Network inputs ($\boldsymbol{I}$) consist of binary mask ($\boldsymbol{S)}$ plus spatially uncorrelated noise; the strength of input is modulated by $\boldsymbol{\omega.}$ (b-c) Prediction 1 (Fig. 1c): Variable modular output patterns from uniform stimulus drive. (b) Example of uniform input drive plus noise, which produces modular output pattern. (c) Different noise inputs produce a variety of modular spatial patterns. (d-f) Prediction 2 (Fig. 1d): Structured stimulus input drive designed to match the characteristic wavelength of the network (d) selectively biases the spatial patterns of output activity, with some variation across trials depending on noise (e). Yellow contour lines show layout of input patterns. (f) Trial-averaged response to structured stimulus input shows strong, specific overlap with input pattern. (g) Correlation of input drive pattern to output activity, for single trial (*left*) or trial-averaged (*right*), compared to trial-shuffled controls (average across 40 simulations for 10 different stimulus input patterns, p<0.01, WSR). (h) The similarity of output activity to its input drive pattern is influenced by the strength of the stimulus input drive (stimulus amplitude used in E-G indicated with star). Our *in vivo* results most closely resemble the low stimulus amplitude condition, suggesting that activity evoked in developing networks by opto-stimulation occurs in noisy conditions.

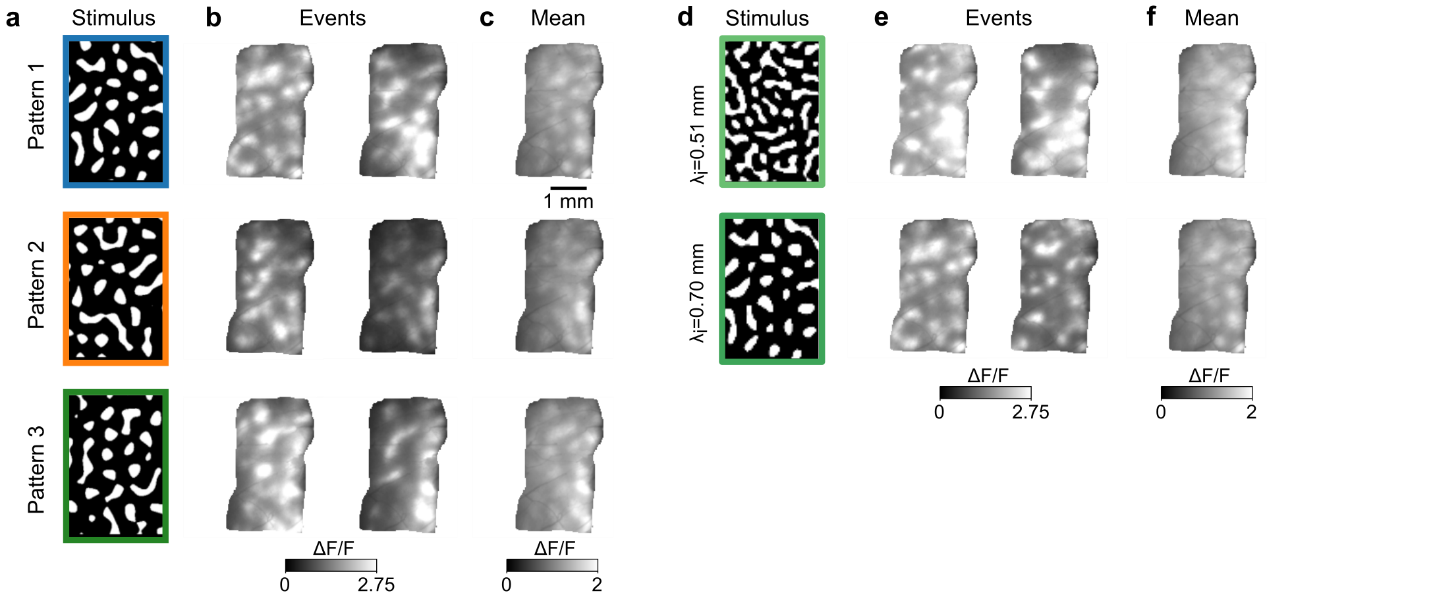

**Fig. S6. Spatially structured opto-evoked activity without spatial filtering.** (**a**) As in Fig. 3a, three example optogenetic stimulation patterns. (**b-c**) Opto-evoked ΔF/F fluorescence activity without spatial filtering, for individual events (**b**) and mean responses across trials (**c**). (**d-f**) same as a-c, except for optogenetic stimuli of differing wavelengths, corresponding to data shown in Fig 4d-f.

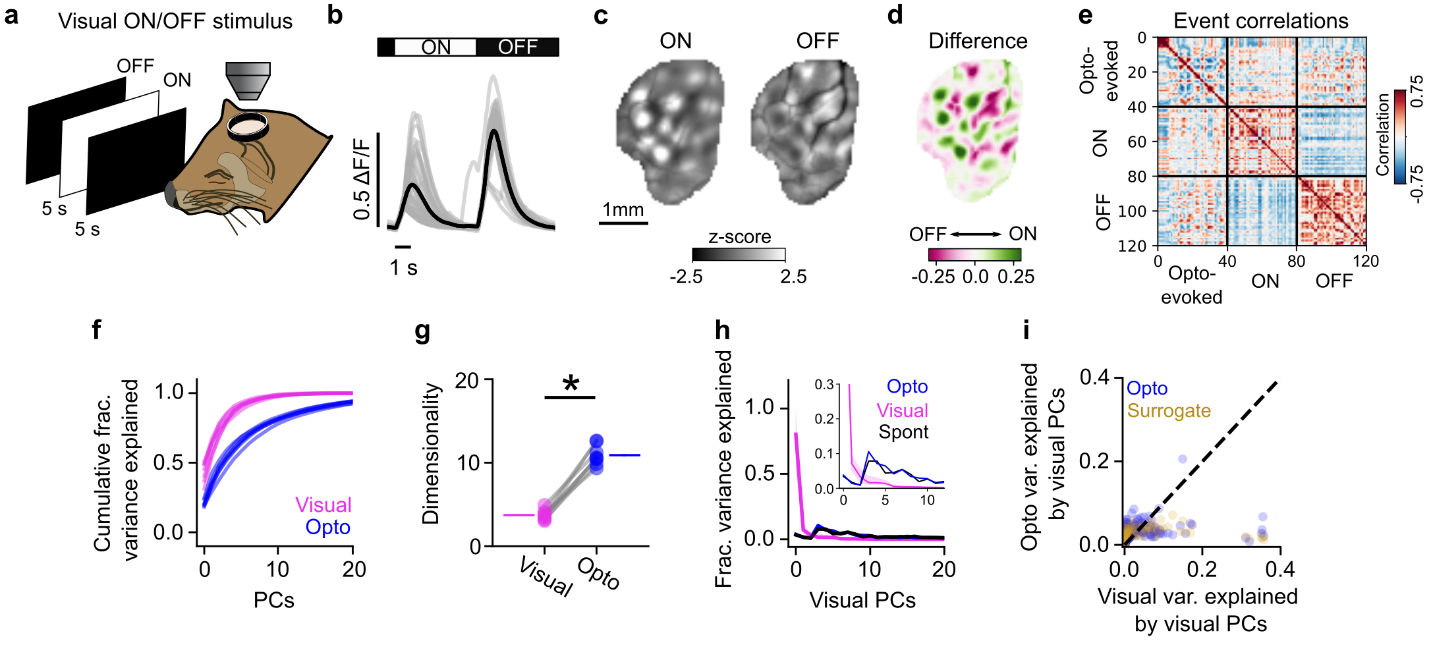

Fig. S7. Luminance evoked visual responses are highly reliable but poorly explain variation in opto-evoked activity. (a) Experimental schematic. (b) Full-screen increases (ON) and decreases (OFF) in luminance drive reliable visually evoked responses through the closed eyelid. (c) Mean ON (left) and OFF (right) responses are modular. (d) Difference map showing ON – OFF selectivity. (e) Trial-to-trial correlation matrix comparing opto-evoked activity and visually-evoked ON and OFF responses. Note reliability within and selectivity across ON and OFF responses, together with weaker and variable similarity to opto-evoked responses. (f-g) Visually-evoked ON/OFF responses are less variable and lower dimensional than opto-evoked responses, requiring fewer principal components to explain the majority of variance in the data (F) and having significantly lower dimensionality (g). (h) Projecting opto-evoked, spontaneous, and cross-validated visual events into the principal component space of visually-evoked activity, the first two PCs explain the majority of the variance of visual activity, while the majority of variance is concentrated on higher PCs for opto and spontaneous events. Inset: enlarged view of first 10 PCs. (i) The PCs of visually-evoked activity that explain the majority of variance of this activity poorly explain the variance in opto-evoked activity, and are indistinguishable from surrogate events (i).

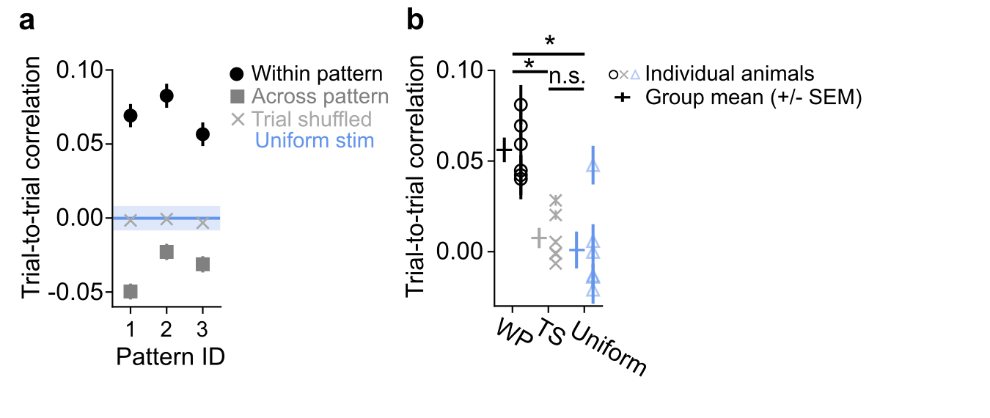

Fig. S8. Structured optogenetic stimulation drives more reliable cortical responses than uniform stimulation. (a) Trial-to-trial correlations between individual trials are greater when comparing trials driven by the same input pattern (within pattern, circles) versus trials driven by non-matching input stimuli (across pattern, squares). Additionally, within pattern trial-to-trial correlations are also greater than both trial-shuffled responses (x’s) and responses to uniform optogenetic stimulation (Blue line, mean trial-to-trial correlation, +/-SEM). Data shown for 1 representative experiment. (b) Across 6 animals, structured optogenetic stimulation (within pattern - WP) drives more reliable responses than trial shuffled (TS) and uniform optogenetic stimulation. Within pattern data are averaged across all 3 stimuli (shown as mean +/- average SEM). Crosses indicate mean (+/- SEM) for each group.

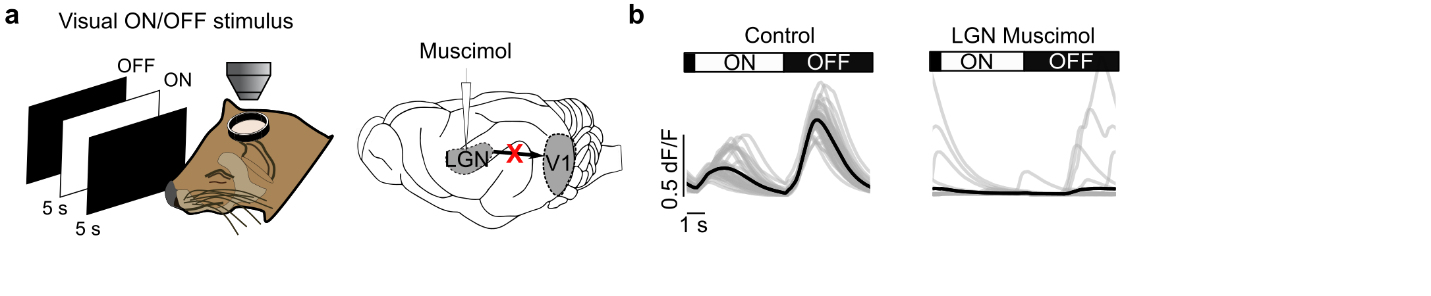

Fig. S9. Pharmacological silencing of feedforward LGN inputs to cortex. (a) Experimental schematic. Feed-forward activity is blocked via muscimol injection to LGN. Silencing is confirmed by measuring responses to full-screen luminance changes. (b) Mean response in V1 to luminance changes prior to (*left*) and following (*right*) muscimol injection confirms silencing of feed-forward projections. Black trace indicates stimulus triggered mean response, grey indicates individual trials (n trials=40). Peaks in activity following LGN muscimol are poorly timed to visual stimulus, and likely reflect spontaneous cortical activity.

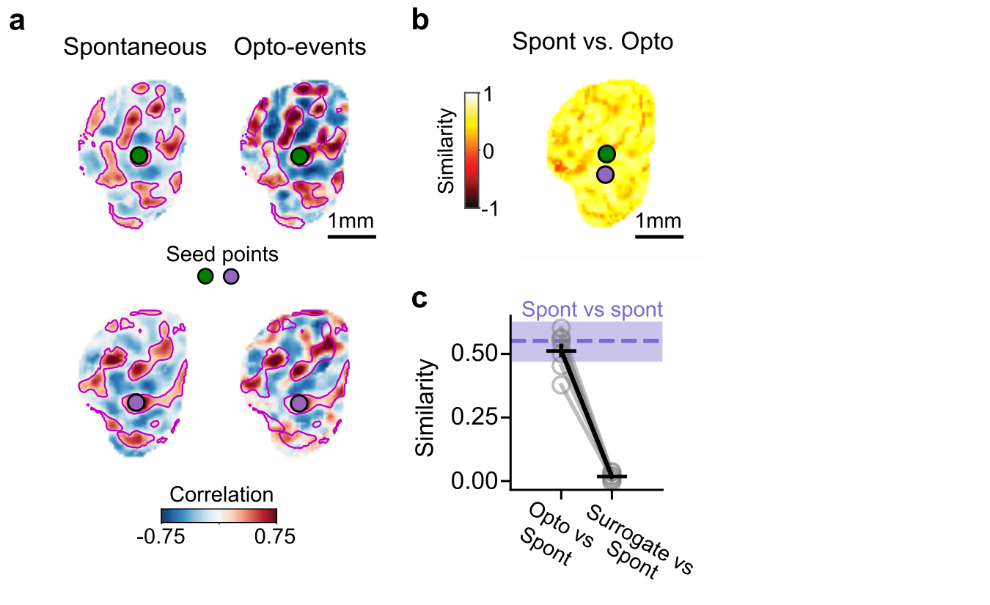

Fig. S10. Uniform opto-evoked events reveal large-scale correlated networks that are highly similar to spontaneous networks. (a) Spatial correlations across spontaneous (*left*) and opto-evoked (*right*) events show highly similar patterns for corresponding seed points. (b) Similarity of spontaneous and opto-evoked correlation networks is high for all seed points. (c) Mean similarity of spontaneous vs opto-evoked networks is equivalent to that for spontaneous activity with itself (purple: spontaneous vs spontaneous, mean (+/- SEM)) and is significantly greater than surrogate data (Mean opto vs spontaneous similarity: 0.512 (+/- 0.028); opto vs surrogate similarity: 0.018 (+/- 0.006), p<0.05, WSR, n=7 animals).

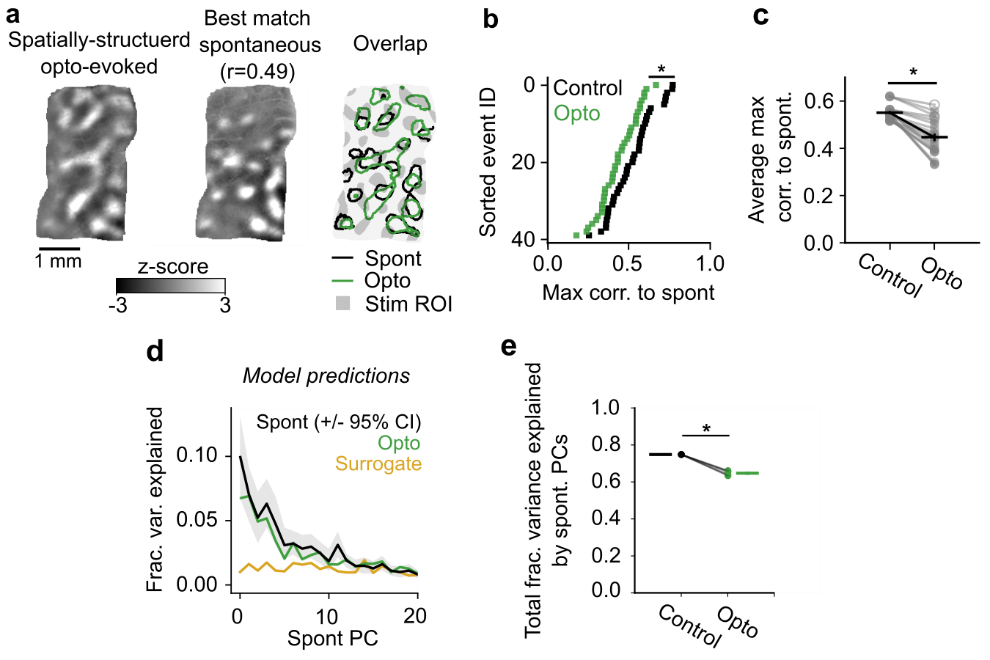

Fig. S11. Overlap of spatially structured optogenetic stimuli to spontaneous activity.

(**a**) Additional example of opto-evoked event (left, green), its maximally correlated spontaneous event (middle, black), and the overlap in active domains on top of the stimulus input pattern (right, grey). Data for stimulus pattern shown in Fig 3. (**b**) Maximum correlation to spontaneous activity for each individual trial. Opto (green) correlations from a single example stimulus pattern shown in (a), Null (black) correlations from random event-matched spontaneous vs spontaneous subsamples (n=500 simulations, see Methods). (**c**) Average maximum correlation across trials for each stimulus pattern (n=21 stimuli, pooled across animals. p<0.001 for 19 of 21 stimulus patterns. Closed circles=stimuli with p<0.001, open circles=n.s). (**d-e**) Computational modeling predictions (h=0.4), showing that structured inputs can drive patterns with novel components that are not entirely explained by spontaneous events. (**d**) Projections of activity patterns driven by spatially structured inputs (Opto, green) onto control activity patterns driven by non-structured noise (Spont, black). Surrogate events (yellow) for *in silico* activity patterns generated the same way as *in vivo* activity patterns (see Methods), so as to maintain statistics of image but randomize spatial pattern. Cross validated control projections done by randomly subsampling event-matched spontaneous patterns and projecting them onto a separate event-matched size spontaneous test distribution, as in Fig 6g. (**e**) Total variance explained by spontaneous PCs, for both control and spatially structured driven datasets (summed across first 40 components). Spontaneous PCs explain a significantly lesser proportion of the variance in the Opto data, indicating that spatially structured stimuli can drive outputs with novel components.

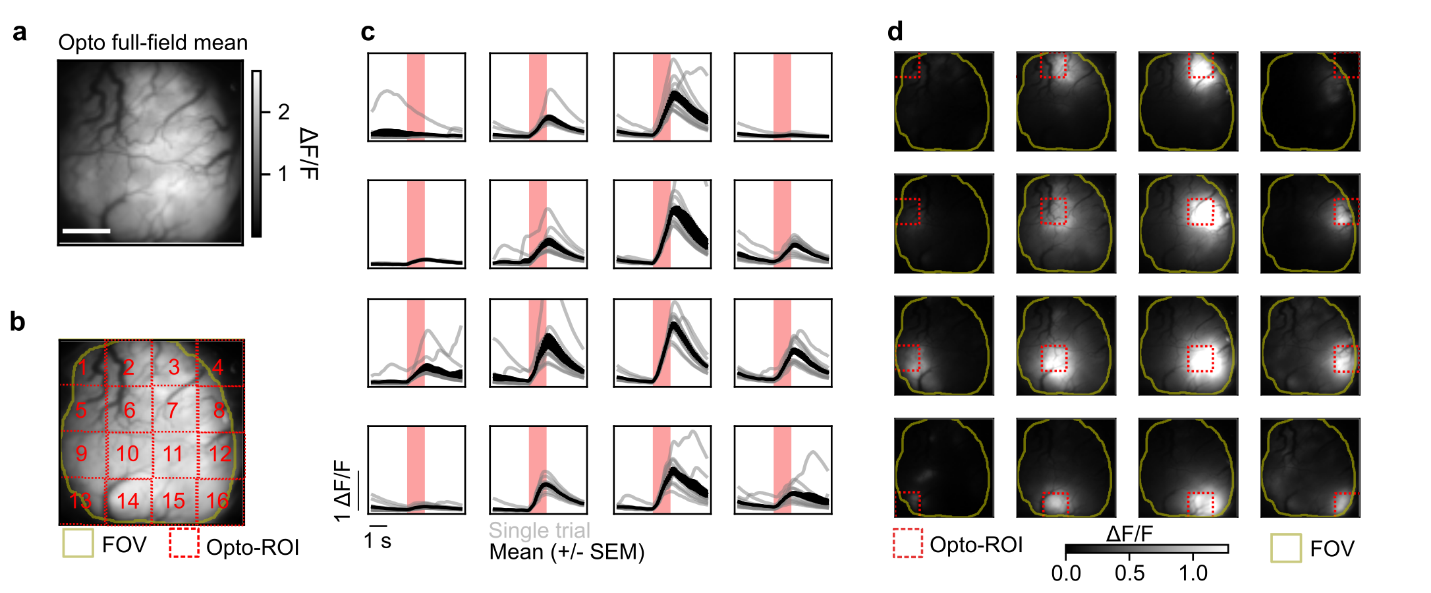

Fig. S12. Response to optogenetic stimulation across imaging field-of-view. (a) Average response to full-field optogenetic stimulation shows broad opto-response across millimeters of FOV. (Scale bar=1 mm). (b) Testing responsiveness to optogenetic stimulation with a 4 x 4 grid stimulus. Each grid square is sequentially stimulated (1 sec duration, 11 trials). (c) Stimulus triggered average response for pixels within stimulus ROI, showing robust and reliable response to optogenetic stimulation across the FOV (grey=individual trials, black=mean (+/- SEM)). ROIs on edge of FOV are poorly driven, due to only partial coverage with stimulus ROI. (d) Mean response to optogenetic stimulation across trials. Dashed red=opto-stimulus ROI, yellow=imaging FOV.

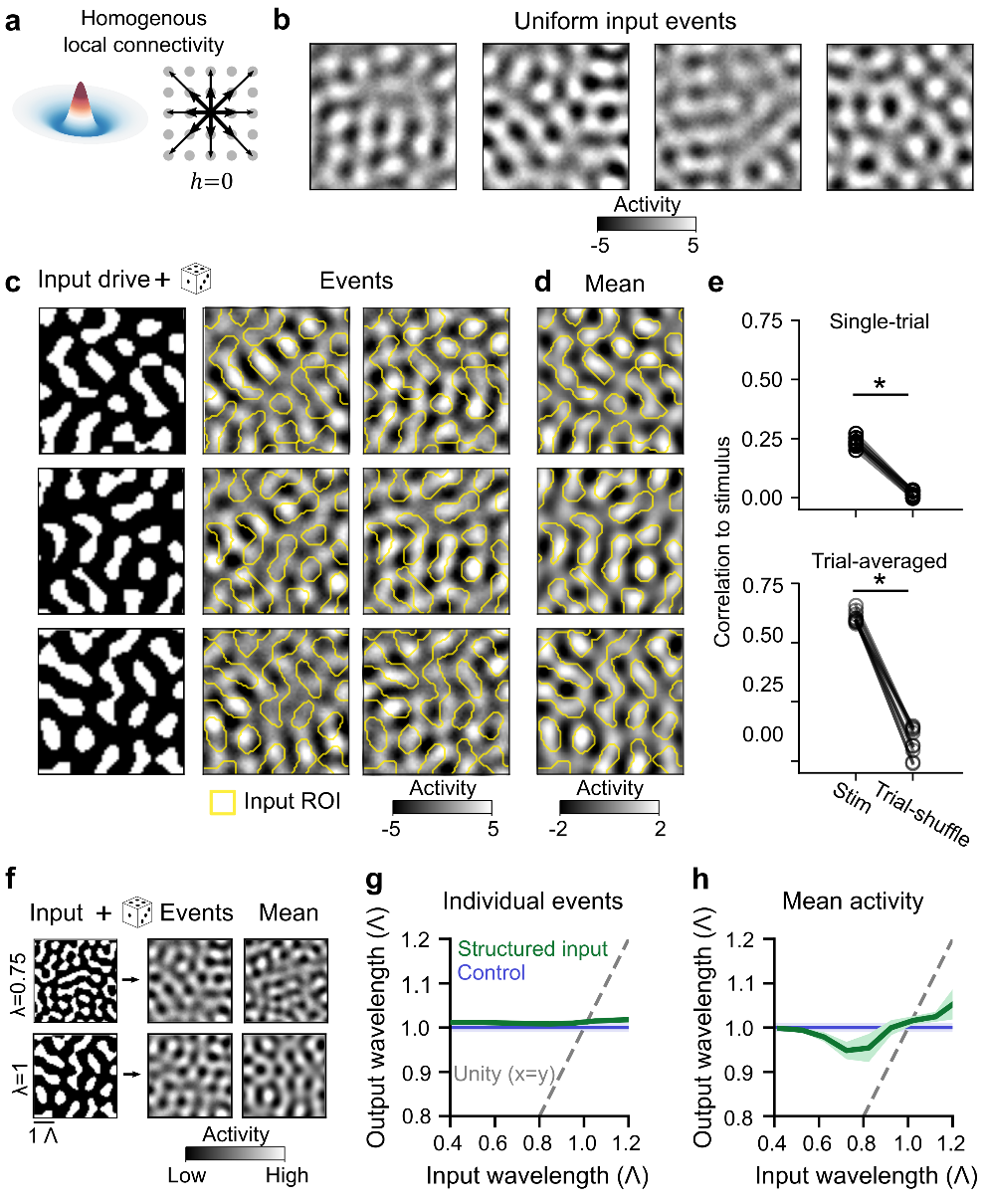

**Fig. S13. Model predictions in a network with homogeneous connectivity.** The predictions in Fig. 1c-e are not dependent on heterogeneity of network connections and are largely unchanged in a model with homogenous local connectivity (compare Fig. S4). (**a-e**) As in Fig. S4a, c, e, f, g, respectively, but for homogenous (i.e. symmetric in rotation and translation, see Methods) cortical interaction. (**f-h**) As in Fig. 4a-c, but for homogenous connectivity.

Movie S1. Uniform optogenetic stimulation of developing V1 reliably evokes modular cortical activity. Modular activity emerges in response to uniform cortical stimulation, with multiple coactive local regions distributed across several millimeters of cortex. Sequential trials of unstructured uniform input reliably elicit modular activity of a consistent spatial wavelength, yet the specific spatial pattern of activity varies greatly across trials. Movie shows time series of 6 trials of uniform optogenetic stimulation. White bar in upper left indicates timing of presentation of optogenetic stimulation. Trace shows mean activity (ΔF/F) within imaging FOV over duration of video; red bands indicate periods of uniform optogenetic stimulation. Images are shown as ΔF/F. Scale bar (lower left): 1 mm.
